# The pig X and Y chromosomes: structure, sequence and evolution

**DOI:** 10.1101/012914

**Authors:** Benjamin M. Skinner, Carole A. Sargent, Carol Churcher, Toby Hunt, Javier Herrero, Jane Loveland, Matt Dunn, Sandra Louzada, Beiyuan Fu, William Chow, James Gilbert, Siobhan Austin-Guest, Kathryn Beal, Denise Carvalho-Silva, William Cheng, Daria Gordon, Darren Grafham, Matt Hardy, Jo Harley, Heidi Hauser, Philip Howden, Kerstin Howe, Kim Lachani, Peter J.I. Ellis, Daniel Kelly, Giselle Kerry, James Kerwin, Bee Ling Ng, Glen Threadgold, Thomas Wileman, Jonathan M D Wood, Fengtang Yang, Jen Harrow, Nabeel A. Affara, Chris Tyler-Smith

## Abstract

We have generated an improved assembly and gene annotation of the pig X chromosome, and a first draft assembly of the pig Y chromosome, by sequencing BAC and fosmid clones, and incorporating information from optical mapping and fibre-FISH. The X chromosome carries 1,014 annotated genes, 689 of which are protein-coding. Gene order closely matches that found in Primates (including humans) and Carnivores (including cats and dogs), which is inferred to be ancestral. Nevertheless, several protein-coding genes present on the human X chromosome were absent from the pig (e.g. the cancer/testis antigen family) or inactive (e.g. *AWAT1*), and 38 pig-specific X-chromosomal genes were annotated, 22 of which were olfactory receptors. The pig Y chromosome assembly focussed on two clusters of male-specific low-copy number genes, separated by an ampliconic region including the *HSFY* gene family, which together make up most of the short arm. Both clusters contain palindromes with high sequence identity, presumably maintained by gene conversion. The long arm of the chromosome is almost entirely repetitive, containing previously characterised sequences. Many of the ancestral X-related genes previously reported in at least one mammalian Y chromosome are represented either as active genes or partial sequences. This sequencing project has allowed us to identify genes - both single copy and amplified - on the pig Y, to compare the pig X and Y chromosomes for homologous sequences, and thereby to reveal mechanisms underlying pig X and Y chromosome evolution.

## Introduction

The therian (marsupial and placental mammal) sex chromosomes evolved originally from a homologous pair of autosomes about 170-180 million years ago (Cortez et al. 2014; Livernois et al. 2011), and have become extensively differentiated in terms of structure and sequence content. The gene content and organisation of the emergent X chromosome has been subject to strong conservation across different mammalian species with retention of much of the ancestral X (Ross et al. 2005; Bellott and Page 2010). In contrast, the acquisition of a dominant male sex-determining function and accumulation of male benefit genes (e.g. genes involved in regulating male germ cell differentiation) on the Y chromosome has been accompanied by (a) the genetic isolation of much of the Y (suppression of recombination with the emergent X), (b) subsequent degeneration and loss of much of the ancestral Y gene content and (c) selection for dosage compensation in XX females to restore equivalence of gene expression between males and females for loci carried on the X that have degenerated or do not have a homologue on the Y (Bachtrog 2013; Graves 2010). Selection has also acted to retain a strictly X-Y homologous pseudoautosomal region (PAR) that permits X-Y pairing during meiosis and within which there is obligate recombination between the sex chromosomes. The gene and sequence content of the PAR varies between mammalian species and this reflects processes of expansion and pruning of the PAR in different mammalian lineages (Otto et al. 2011).

Comparisons of X chromosome sequences from several mammalian species have confirmed strong conservation of gene sequence and order (Chinwalla et al. 2002; Sandstedt and Tucker 2004). Groenen et al. (2012) published the first assembly of the porcine X chromosome as part of the initial description of the pig genome sequence, and again this demonstrated conservation of synteny across the chromosome. Nonetheless, sequence gaps and ambiguities remained within this first assembly, complicating genomic studies within pigs, and comparative studies between mammals.

In contrast to the broadly conserved X chromosomes, the hemizygous nature of the Y chromosome and suppression of recombination, in combination with normal processes of genome evolution, have led to a gradual degeneration of the chromosome over time, a large number of rearrangements, and colonisation by sequences from the X and autosomal chromosomes. Newly introduced genes will either drift or degenerate, or selection may act on variants to fix new genetic functions on the Y, particularly where these confer a benefit to the male. The haploid state of the sex chromosomes in males has generally led to the accumulation of male gametogenesis genes on both X and Y chromosomes (Vallender and Lahn 2004). A further consequence of the non-recombining status of the differential region of the Y is the relaxation of restraint on sequence amplification, leading to generation of ampliconic regions containing amplified gene and sequence families (Bellott and Page 2010).

The highly repetitive nature of many regions of mammalian Y chromosomes has impeded the generation of complete chromosome sequences; while there are tens of mammalian genomes sequenced, only a small fraction have a Y assembly. These few assemblies, plus several partial sequence assemblies have permitted the elucidation of chromosome topology and gene order in human and chimpanzee (Hughes et al. 2010), mouse (Soh et al. 2014), cattle (Elsik et al. 2009), horse (Paria et al. 2011), cat and dog (Li et al. 2013a). These works clearly show the great divergence in gene content, order, structure and sequence between Y chromosomes from different mammalian species. However, little data is available on the porcine Y chromosome sequence, gene order and their relationship to the X, despite the recent sequencing project for the pig genome (Groenen et al. 2012). Much of our knowledge of Y gene order comes from Quilter et al. (2002) who combined radiation hybrid mapping data with physical mapping of BAC clones to generate an ordered gene list.

This paper presents a second-generation, much improved assembly and gene annotation of the porcine X chromosome. We present also a body of Y chromosome sequence, which has permitted a first-generation assessment of the Y chromosome short arm gene content and order, how this compares to other mammalian Y chromosomes, the evolutionary processes leading to the current Y organisation, and the structural relationships between the porcine sex chromosomes.

## Results

### A second-generation porcine X chromosome assembly

#### Organisation and evolution of the X chromosome

The pseudoautosomal region (PAR) in pig is of a similar size to the PAR in other closely-related mammals (e.g. cattle). We have previously discussed its gene content, its comparison to other mammalian species and the delineation of the boundary region (Skinner et al. 2013). Recently the precise location of the PAR boundary was confirmed to be within the gene *SHROOM2* (Das et al. 2013). The X chromosome assembly (http://vega.sanger.ac.uk/Sus_scrofa/Location/Chromosome?r=X-WTSI) comprises 129,927,919bp of sequence in 5 contigs, with 13 gaps and an N50 length of 4,824,757bp. Compared with the previous 10.2 build, many gaps have been filled and the order of sequences on the chromosome has been updated. Much of this improvement was aided by the use of optical mapping techniques, which helped resolve some of the more repetitive regions of the chromosome such as around the centromere; an example can be seen in the short arm clone CH242-202P13 (see Supplementary Methods for details on how the optical mapping approach was used here). Supplementary figure S1 shows a dot-plot alignment of the 10.2 X with our X assembly, highlighting the regions of the chromosome for which the sequence order has been corrected.

We aligned the X (and available Y) chromosome sequences of nine mammalian species (Figure 1). The previously documented high level of conservation of X chromosome synteny is more apparent with the new pig X, as many of the reported breakpoints from cross-species comparisons to the build 10.2 X were due to errors that have been resolved in the new assembly. Specific lineages though, such as the rodents, have higher levels of chromosomal rearrangement. Li et al. (2013b) produced a genome assembly of a female Tibetan wild boar, and reported regions of the genome with apparent inversions with respect to the Duroc assembly. We compared the X inversions with our new assembly, and found that they lie outside the regions that have changed orientation from the 10.2 X. That is, these remain as potential inversions between Duroc and Tibetan wild boar.

**Figure 1.**
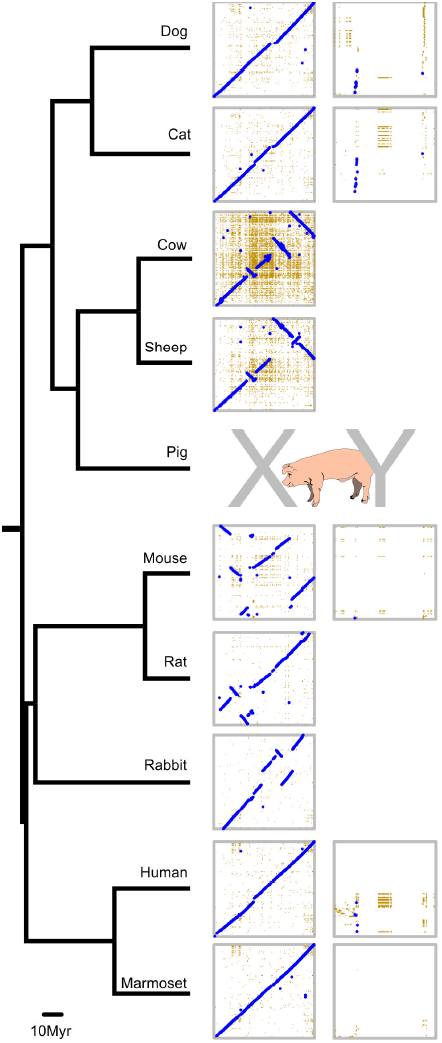
Comparative X and Y map. Sequenced X chromosomes from nine mammals, plus Y chromosomes from five of these, aligned to our pig X and Y assemblies. In each dotplot, the pig chromosome is on the horizontal axis, and the subject chromosome is on the vertical axis. Note the cow sequence is plotted in reverse orientation. High-stringency alignments are shown in blue, with less stringent alignments in yellow. Many lineages (e.g. human, marmoset, cat, dog) retain the ancestral X arrangement. Other lineages (e.g. sheep and cattle) have had small number of rearrangements, while others still (e.g. rodents, rabbit) have a much greater rate of chromosomal change. The Y chromosomes, in contrast, have a highly variable organisation, and none likely resemble the ancestral Y.

#### Gene content of the X

Since the reference assembly of the X chromosome was high quality, the sequence underwent manual annotation, since the chromosome contains complex duplicated gene families such as olfactory receptors along with pseudogenes, which are hard to discriminate using only automatic pipelines. Table 1 shows the updated annotation compared to the 10.2 X build, plus comparable statistics for the annotation of the Y chromosome. The full gene annotation is provided in Supplementary Table S7 and is available through Vega browser. The majority (76%) of annotated loci in pig are shared with human. Again, the improved sequence assembly facilitated by optical mapping analysis assisted the new gene annotation, and many genes from the previous build that were disrupted by gaps have now been completed. The number of long non-coding RNA loci that have been identified on the new assembly has increased from 33 to 84, and, although the functionality of this category of loci is still under debate (Young and Ponting 2013), more and more evidence is coming to light suggesting that at least some of these loci have a functional role within the genome; for example, the lncRNA FIRRE has recently been identified in mouse and human and implicated in having a role in interfacing and modulating nuclear architecture across chromosomes (Hacisuleyman et al. 2014). Despite very low levels of sequence conservation between species, we find evidence for a *FIRRE*-like locus in the syntenic region of pig (OTTSUSG00000005757).

**Table 1.** Comparison of gene content between porcine build 10.2 X, our updated build of the X(X-17), and the new Y annotation (Y-12). The number of identified coding genes has been substantially increased, and brings the porcine X closer in gene content to other well-sequenced mammalian X chromosomes (e.g. human and mouse).

|  | 10.2X | X-17 (WTSI-X) | Y-12 (WTSI-Y) |
| --- | --- | --- | --- |
| **Total Number of Genes** | 632 | 1031 | 196 |
| **Total Number of Protein Coding** | 422 | 687 | 75 |
| Known Protein Coding | 211 | 435 | 18 |
| Novel Protein Coding | 210 | 244 | 57 |
| Putative Protein Coding | 1 | 8 | 0 |
| **Non-coding genes** |  |  |  |
| LincRNA | 20 | 62 | 11 |
| Antisense | 11 | 22 | 0 |
| Sense intronic | 0 | 1 | 1 |
| **Pseudogenes** |  |  |  |
| Processed Pseudogenes | 140 | 196 | 29 |
| Unprocessed Pseudogenes | 2 | 21 | 72 |
| Unitary Pseudogenes | 3 | 11 | 0 |
| Transcribed Unprocessed Pseudogenes | 1 | 3 | 3 |
| Transcribed processed Pseudogenes | 0 | 0 | 0 |

**Table 2.** Genes on pig Yp with expression status and supporting evidence. A morecomprehensive table may be found in the supplementary data (table S3). \**ZFY*: we have resequenced and have no evidence for the frameshift in the accessioned version of the transcript. Our cDNA data agree with the Y fosmid sequences, suggesting that the Y ORF is intact.

| Gene (*alias*) | Status in our sequence | Expression | Reference |
| --- | --- | --- | --- |
| *KDM6A (UTY)* | Incomplete in build. ORF covering just over 50% wrt X. | Ubiquitous | Quilter et al. 2002 |
| *DDX3Y (DBY)* | Complete | Ubiquitous | Quilter et al. 2002 |
| *ZRSR2* | 5' end incomplete and inverted wrt main body of exonic material | n/a |  |
| *AP1S2* | Single exon fragment | n/a |  |
| *USP9Y*<br>(*DFFRY*) | Complete | Ubiquitous | Quilter et al. 2002 |
| *USP9Y*<br>(*DFFRY*) | Partial copy: not processed | n/a |  |
| *EIF1AY* | Complete | Ubiquitous | Quilter et al. 2002 |
| *RPL5L* | Partial copy: processed | n/a |  |
| *MBTPS2* | Partial gene fragment | n/a |  |
| *EIF2S3Y* | complete | Ubiquitous | Quilter et al. 2002 |
| *ZFY\** | Complete | Ubiquitous? | Quilter et al. 2002 |
| *SEPHS1* | Processed complete ORF | n/a |  |
| *HMGB1* | Processed complete ORF | n/a |  |
| *AMELY* | Complete | Tooth development | Hu et al. 1996 |
| *TMS4BY* | Complete | Multiple tissues in EST database |  |
| *TRAPPC2Y* | Partial, processed pseudogene | Intestine, prostate in putative Y-derived ESTs | This study |
| *OFD1Y* | Complete | Here: testis, kidney, brain<br>EST support: intestine, spleen, adrenal, adipose | This study |
| *GPM6B* | Not processed. Disrupted - two fragments | n/a |  |
| *KDM5D*<br>(*SMCY*) | Exons present, but no good evidence of complete ORF | Spliced transcript detected | Quilter et al. 2002 |
| *BTG1* | Processed. No ORF. | n/a |  |
| *BCOR* (*ANOP2*<br>*MAA2*<br>*MCOPS2*) | Not processed. Missing 5' end. Internal exon matches also close to *TXLNGY* | n/a |  |
| *TXLNGY* | Complete | Not restricted: liver, brain, kidney, testis | This study |
| *UBA1Y*<br>(*UBE1Y*) | Potential for large ORF | Ubiquitous | Quilter et al. 2002 |
| *CUL4BY* | Potential for ORF. cDNA completed. | Testis, kidney, brain. High in testis, low in other two tissues | This study |
| *SRY* | Complete ORF | Testis |  |
| *RBMY* |  | Testis ESTs only |  |
| *HSFY* | Multicopy | Testis, low kidney | Skinner et al, in submission |
| *TSPY* | No perfect match to current Y clones, but not autosomal or X | Probable Y expressed locus; not in fosmids characterised |  |

Some genes with updated annotation from build 10.2 stand out as being of particular biological interest. Comparing the human and pig X chromosomes there are 38 coding loci in pig that are not found in human. Twenty-two of these coding genes, plus five novel pseudogenes, are in olfactory gene clusters. Pigs are known to have a large OR repertoire (Nguyen et al. 2012), and this adds to the reported collection. Figure 2 shows the improved assembly and annotation around one of the olfactory region clusters on Xq. The region lay within an inversion in the 10.2 assembly, corrected here and matching the gene order on the human X. The full list of genes present on the pig X chromosome, but not on the human X is provided in supplementary table S9.

**Figure 2.**
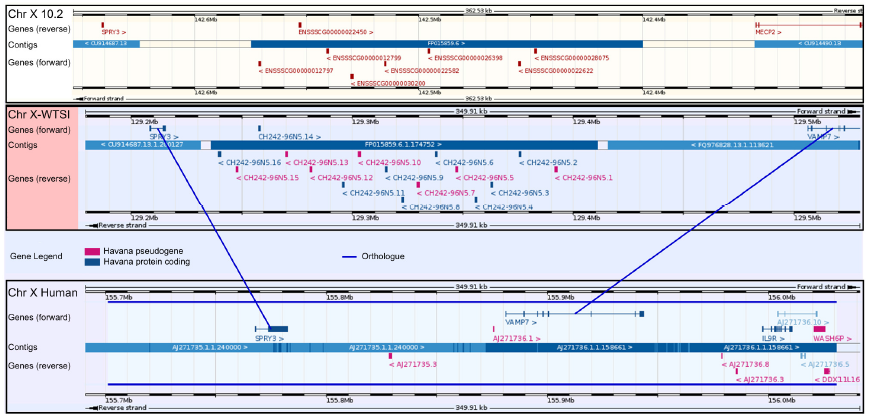
Assembly improvement around an olfactory receptor gene cluster expansion. Pigs have an expansion of olfactory receptors throughout the genome; two clusters lie on the pig X, not found in humans. The region provides an example of the sequence improvement from optical mapping between the 10.2 assembly (top; image from Ensembl) and the current assembly (centre), with a comparison to the corresponding region on the human X (bottom; both images from Vega). The improvement in the current assembly allowed reorientation (Ensembl image has been flipped for clarity), the correct positioning of the VAMP7 gene which was previously annotated on an unplaced scaffold and improved annotation of the genes (blue) and pseudogenes (pink) within the region.

Besides the additional genes compared to human, there are also 11 protein coding genes which are present on the human X, which have been annotated as unitary pseudogenes (also known as Loss Of Function genes) on the pig X (Supplementary Table S10). These include *GUCY2F* (OTTSUSG00000005153) which in humans has been suggested as a possible candidate for involvement in X-linked retinitis pigmentosa (RP) (Yang et al. 1996). Pig models are now being developed for studying RP, taking advantage of the similarities between human and porcine retinal development (e.g. Ross et al. 2012; Fernandez de Castro et al. 2014). These models benefit from an improved understanding of the status of orthologues of disease-related genes:

*AWAT1* (OTTSUSG00000002936) is another unitary pseudogene. This gene is found on the X chromosome in a wide range of mammals, including cattle, humans and opossums (Holmes 2010). It encodes an acyl CoA wax alcohol acyltransferase involved in sebum production, and in humans is expressed in sebocytes, aiding in the prevention of surface desiccation of the skin (Turkish et al. 2005). The gene is a member of a family including *DGAT1* and *DGAT2*, which have previously been associated with backfat thickness and intramuscular fat content respectively in pig breeds (Cui et al. 2011). It is likely that this pseudogenised *AWAT1* in Duroc pigs represents a species-wide trait, as no transcripts for *AWAT1* appear in the EST databases from any pig breeds.

Other identified unitary pseudogenes include *ITIH6*, a trypsin inhibitor, and *RAB4*, a member of the RAS oncogene family, as well as a number of zinc finger proteins and a transmembrane protein. Whether some of these loci were ever functional in pig, or merely reflect conserved regions that became functional at some point in the human lineage, is open for debate.

Other regions of difference lie in the cancer/testis (*CT*) antigen clusters found in humans and other primates but which are significantly reduced n pig. This is in line with evidence that enlarged primate CT antigen clusters arose due to a recent amplification in primates (Zhang and Su 2014), perhaps driven by a retrotransposition event. Their potential functions remain unknown, though they may have been involved in primate speciation.

Lastly it was found that *INE1* (Inactivation-escape-1) is not present in pig. This is a non-coding transcript with unknown function within an intron of *UBE1X* (Thiselton et al. 2002). It appears to be unique to humans, where it escapes X inactivation in females.

### A first generation porcine Y chromosome sequence assembly

Successfully assembled repetitive portions of Y chromosomes have been generated only for a limited number of species (see for example the human, mouse or chimpanzee Y; Hughes et al. 2010; Soh et al. 2014). Given the highly repetitive nature of the long arm of the pig Y chromosome (Quilter et al. 2002; Skinner et al. 2013), we targeted the short arm, which contains most, if not all, of the single-copy material.

The strategy employed was to prepare fosmid clones from flow-sorted Y chromosomes (origin Duroc) and initially to use fingerprint analysis to establish a framework of clones and to identify those inserts belonging to highly repetitive regions. Previously identified BAC clones (origin Meishan, from a Large White female/Meishan male cross: Anderson et al. 2000) containing Y chromosome genes conserved between several mammalian species served as anchor points for the elaboration of more complete sequence contigs. All of these BAC clones are known to map to the short arm of the Y (Quilter et al. 2002). Overlaps between BAC and fosmid clones were identified to extend the contigs, and selected clones within each contig were used for physical mapping (by metaphase-, interphase-and fibre-FISH). This, combined with existing data from linkage and radiation hybrid mapping, allowed us to produce an ordered map of the genes and contigs on the short arm of the Y. Selected repeat-containing clones were also sequenced to sample the diversity of repetitive sequence content of the Y chromosome. The final contigs we are predominantly of Duroc origin, with small regions of Meishan where no Duroc sequence was available. Supplementary table S1 summarises the ordered clones that we were able to incorporate into contigs including small contigs that could not be anchored to the physical map. The resulting male-specific Y sequence data is available at http://vega.sanger.ac.uk/Sus_scrofa/Location/Chromosome?r=Y-WTSI A two-pronged strategy was used to identify potential coding sequences in the body of fosmid sequence data. First, all fosmid sequence was subjected to gene prediction algorithms and potential coding sequences compared to the non-redundant protein database. Second, extensive BLAST analysis of known mammalian Y-linked genes and porcine X-linked homologues against the Y fosmid sequence dataset was used to identify porcine homologous sequences. The established intron/exon structure of these genes was used to drive the assembly of more extensive coverage of the porcine Y homologue. These sequences were used to identify ESTs in order to confirm expression status. Select genes were further tested by RT-PCR in a range of tissues.

### Organisation of the porcine Y chromosome

Figure 3 summarises the genes identified on the Y and their order, and the distribution of families of mammalian repetitive sequences across the short arm. The relative order and orientation of genes on the differential region of the Y short arm was established using a combination of the assembled contig sequences and high-resolution FISH with DNA fibres prepared by the Molecular Combing approach. The fibre-FISH also allowed an estimation of the gap sizes between adjacent contigs (see supplementary figure S2 for detailed presentation of supporting evidence). The genes are organised into two main blocks of low copy number sequences. These two blocks are separated by a region containing highly amplified sequences including the *HSFY* gene family (see Skinner et al, in submission). This amplified block maps to cytogenetic band Yp1.2 and is estimated from FISH analysis to be about 5Mb in length. Our final mapped sequence comprises seven contigs in the distal block and seven contigs in the proximal block. The final contig sequences were analysed for repeat content with RepeatMasker, and the densities of repetitive content found are shown in the column to the left of Figure 3. Two large (~400kb) contigs were assembled based on sequence overlaps and confirmed by fibre-FISH, but could not be assigned to the physical map (supplementary figure S3). These unplaced contigs did not contain any identifiable genes, and are included in a separate assembly in VEGA.

**Figure 3.**
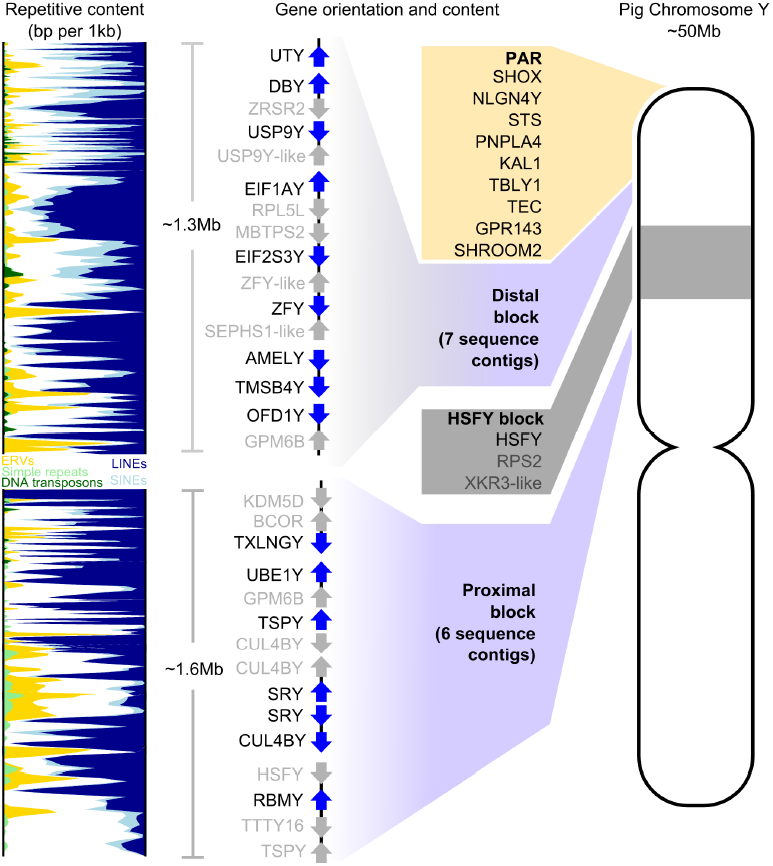
The pig Y chromosome. The organisation of the pig Y chromosome. All identified male-specific single copy genes are on the short arm and split into two blocks by the ampliconic *HSFY* region. The genes (blue) and pseudogenes (grey) are shown within each block. To the left of the figure is the density of repetitive content based on a 10kb sliding window within the sequence contigs we obtained.

Very few low-copy loci have been identified on the Y long arm. Indeed, FISH with clones containing male-specific repeats paints the entire long arm, indicating that it is mostly composed of repetitive sequences (e.g. Quilter et al. 2002). Although we attempted to sequence one of these clones, it was not possible to assemble a framework physical or sequence map from the repeats it contained, as most sequences collapsed into a single contig. The sequences we obtained from this clone are shown in the supplementary data and belong to previously published male-specific pig repeat families (Mileham et al. 1988; McGraw et al. 1988; Thomsen et al. 1992; Pérez-Pérez and Barragán 1998). Metaphase FISH did, however, reveal a small low-copy region at the q terminus (see supplementary figure S13).

### Gene-related content of the Y chromosome

The Y chromosome sequence was run through the Otterlace/Zmap analysis pipeline which performs homology searches, *de novo* sequence analysis and gene predictions (Loveland et al. 2012). Repeat masking proved challenging due to the paucity of known pig-specific repeats. Manual annotation resolved this as homology analysis is routinely run on-the-fly within the annotation tools, without repeat masking, to more accurately elucidate gene structures, especially using known Y chromosomal genes from other species as targets, and identifying the pig homologues where present. Many of the ancestral-X-related genes previously reported in other mammalian Y chromosomes are represented either as active genes or partial sequences, some of the latter with supporting ESTs. There is also evidence for rearrangement of a number of Y-linked genes which may have rendered them non-functional or modified their transcription products. RT-PCR analysis for several of these loci across a range of porcine tissues was performed to establish their transcriptional status. A list of the gene loci we identified in this and previous work is given in table 1. A fuller table including genes not tested here but with EST support is provided as supplementary table S3.

### Ampliconic gene sequences

Although our sequence contigs are limited to the low copy regions of the chromosome, the FISH data show regions where gene-containing clones are present in multiple copies. Examples are the *CUL4B* genes, one of which is present as a partial copy in fosmid WTSI_1061-13A2. Probes against this sequence bind multiple targets proximal to *SRY* (and likely proximal to *RBMY*) as well as additional sites towards the centromeric end of the Y short arm (see supplementary figure S13). This indicates that *CUL4B* is likely to be an ampliconic gene. The sequence supports gene expression from a ‘full-length’ version of the sequence centromeric to *SRY*, and RT-PCR shows expression in testis, kidney and brain. A similar situation exists for clones containing *TSPY*.

### Regions of X-Y Homology

We examined all the sequenced Y clones for homology with the X. Outside the PAR, two large (~50kb) regions of XY homology were seen, the remaining shorter regions comprising individual XY homologous genes and various classes of repetitive sequences. The overview X-Y comparison is given in supplementary figure S4). Most gene-related sequences show homology to the X.

We examined rates of synonymous substitution in open reading frames between the X and Y copies of genes conserved on Y chromosomes across mammalian species but within different evolutionary strata as defined for the human X and Y (Lahn and Page 1999) (Table 3). Gene pairs have similar substitution rates to their orthologues in other mammals.

**Table 3.** Synonymous substitution rates (K_S_) across Y gene coding regions. Genes from more ancient evolutionary strata as defined in the human X (Lahn and Page 1999) have higher number of synonymous substitutions per synonymous site. The low K_S_ of the *ZFX/Y* genes is consistent with previous description of ongoing gene conversion in other mammalian species (e.g. Slattery et al. 2000; Pamilo and Bianchi 1993).

| Gene pair | Aligned length | $K_S$ | $K_A$ | $K_A/K_S$ | Stratum |
| --- | --- | --- | --- | --- | --- |
| *RBMX/Y* | 1185 | 0.826 | 0.112 | 0.136 | 1 |
| *UBA1X/Y* | 3177 | 0.686 | 0.066 | 0.096 | 2 |
| *EIF1AX/Y* | 435 | 0.568 | 0.006 | 0.011 | 3 |
| *ZFX/Y* | 2406 | 0.157 | 0.016 | 0.102 | 3 |
| *DBX/Y* | 1989 | 0.478 | 0.020 | 0.042 | 3 |
| *AMELX/Y* | 570 | 0.046 | 0.009 | 0.196 | 4 |
| *OFD1X/Y* | 3003 | 0.090 | 0.039 | 0.433 | X-transposed |

A ~50kb region on the distal block encompassing the genes or gene fragments *TRAPPC2-OFD1Y-GPM6B* has high sequence identity to the X even in intronic and intergenic regions, suggesting a possible X-Y transposition (supplementary figure S6). *OFD1Y* is expressed highly in testis with lower expression in kidney and brain (supplementary figure S8), but there is no evidence for expression of the Y copies of *TRAPPC2* or *GPM6B*. Furthermore, *GPM6B* is missing exons 1-3. *GPM6B* exon 1 lies near *SMCY*, possibly indicating an ancient inversion or other rearrangement. Recurrent X to Y transpositions of this region have been suggested before in cattle and dog (Li et al. 2013a; Chang et al. 2013); a gene tree constructed from dog, cattle, pig and horse *OFD1X* and *Y* sequences shows that the pig Y copy is more similar to the *OFD1X* than to any other species Y copies (supplementary figure S7), supporting the idea that this region arose on pig Y via a transposition from the X. A further ~33kb region of homology within the PAR (WTSI-X:1,840,693-1,874,130) can also be found on Xq (WTSI-X:114,853,327-114,886,764), and potentially represents a region of duplication and transposition from the PAR onto Xq.

### Evolution of the porcine Y chromosome

#### Inverted and duplicated blocks of sequence

Inversion and duplication to form palindromes has occurred around both *SRY* and *CUL4BY* (Figure 4), evidenced by sequence and fibre-FISH data for fosmids. The *SRY* gene itself is present in two head to head copies, validated by fibre-FISH experiments using the gene sequence directly as a probe. There are unlikely to be more than two *SRY* copies on the chromosome (qPCR - supplementary figure S10). In both palindromes, the ancestral arm of the palindrome was the proximal copy, and the derived copy is distal. This is evidenced by a disrupted ancestral ERV element outside the *SRY* inversion, and a partially duplicated LINE element overlapping the *CUL4BY* inversion (see supplementary figure S9). The pattern of markers at the breakpoint regions reveals that the *SRY* duplication preceded the *CUL4B* duplication (see Figure 4 and supplementary figure S9). The transposable elements at both inversion boundaries are annotated as specific to the *Sus* lineage, suggesting these are relatively recent duplications. The copy number of *SRY* and *CUL4B* in closely-related suid species is therefore uncertain. The arms of the palindrome have high levels of sequence identity; we could not detect a difference in the *SRY* sequence from clones on one arm versus the other arm. Our sequence contigs do not cover the centres of the palindromes (about 20kb is missing in each), so we do not know if the arms abut - it is possible that there is a short stretch of unique sequence between the arms.

**Figure 4.**
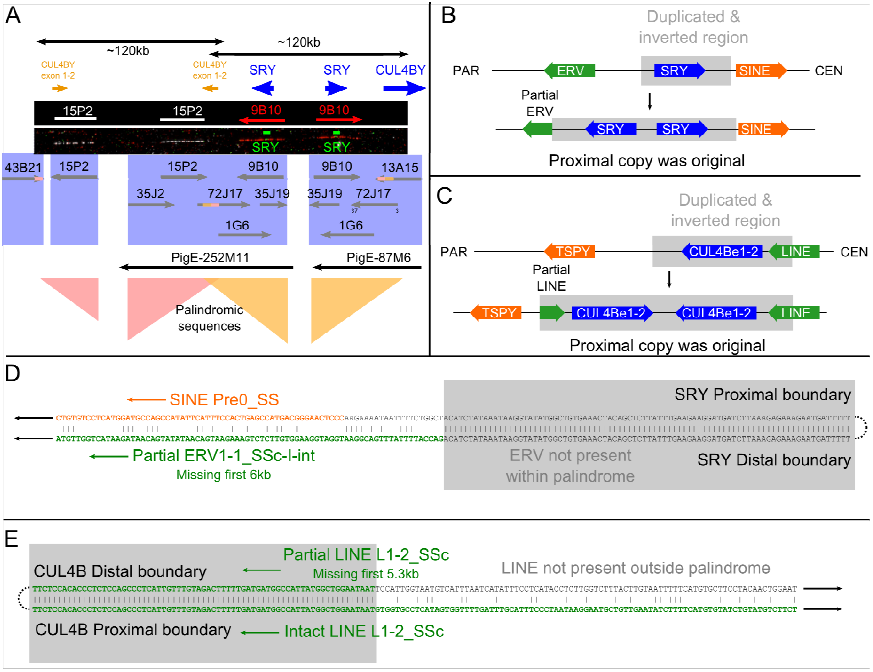
The pig SRY region. The Yp proximal block of genes contains two overlapping palindromes of about 120kb each. These surround the duplicated sequences *CUL4BY* exons 1-2 and SRY. **A)** FISH results from Y fosmid clones and probes for the *SRY* gene are shown with the BAC and fosmid clone sequences found mapping to the region. The inversion boundaries are both identifiable; the *CUL4BY* inversion runs from the last 3kb of 43B21 to within 72J17; the *SRY* inversion begins also within 72J17 and runs to 13A15. A schematic view is also shown of the regions surrounding **B)** the *SRY* and **C)** the *CUL4B* duplications. The *SRY* duplication disrupts and ERV element, revealing the proximal copy to be ancestral. The *CUL4B* duplication copies part of a LINE element, again revealing the proximal copy to be ancestral. The sequence alignments across the inversion breakpoints are shown in more detail for **D)** *SRY* and **E)** *CUL4B*. The order of events was therefore a duplication of SRY including the first two exons of CUL4B, followed by duplication of the region around the *CUL4B* copy. See also Figure S1 for broader context of the region.

A further palindromic region lies at the proximal end of *USP9Y*, covering approximately 50kb, including the final exons 18-43 of the gene. The breakpoints lie well within known transposable elements; one end within a LINE, and the other end within a SINE, and these elements likely facilitated the original rearrangement (supplementary figure S11). Compared to *SRY* and *CUL4BY*, the breakpoints are less well defined, with sequence identity decreasing over some tens of base pairs; this probably reflects a more ancient event than *SRY* and *CUL4BY*.

#### Transposition within the Y chromosome

A distinctive pattern of repetitive elements covering 5kb is found in two Y contigs with 99% sequence identity. This likely resulted from a transposition of sequence at end ERV elements. The sequence originated near *SMCY*, and transposed to *OFD1*; a cluster of ERVs can be seen at the distal end of *OFD1Y* in the distal gene block, and a further ERV cluster distal to *TXLNGY* in the proximal gene block (visible in Figure 3, higher resolution in Figure S2). The breakpoint regions are clearly visible within one ERV element (ERV1-4_SSC-I-int) upon alignment (see supplementary figure S5), while the breakpoint at the *OFD1Y* distal end has been further disrupted by an L1 LINE element insertion. These repetitive elements are all annotated as being *Sus* lineage specific, and show evidence for recent dynamic activity of transposable elements on the Y chromosome.

#### Structural rearrangements compared to other species

We generated likely pathways of gene-only rearrangements from the ancestral Y chromosome to pig, using data from (Li et al. 2013a) updated with pig and cattle (Elsik et al. 2009) Y chromosomes (supplementary figure S12). In pig, as in other species, the *USP9Y-DBY-UTY* genes are the only cluster to retain proximity from the ancestral Y chromosome. Global alignments of chromosome content with other available Y chromosome assemblies are also shown in Figure 1. This shows the striking rate of Y-chromosomal change across mammals. Little of the Y sequences align, save for the genic regions, and these show gross levels of rearrangement. In particular, the comparison highlights some of the ancestral gene families that have become amplified in different lineages, e.g. *HSFY* in pigs and *TSPY* in dogs.

## Discussion

We present here an updated and substantially improved assembly of the pig X chromosome, and a first-generation sequence and map of the pig Y chromosome with expression analysis of many of the genes it contains. This sequence has also allowed us to recover information on the evolution of the Y chromosome, and how this relates to sex chromosome evolution in other mammals.

### An updated assembly and annotation of the porcine X chromosome

The picture of mammalian X chromosomes is one of general structural stability; see Figure 1. Specific lineages, such as rodents, have many more X rearrangements than the others, but these species are characterised by a globally higher number of chromosomal rearrangements (Stanyon et al. 1999). Apparent inversions and translocations in the pig X, relative to the ancestral X, detected in previous builds are corrected here to an order more reflective of the inferred ancestral X chromosome. Similar findings may be seen with other mammalian X chromosomes as the quality of the assemblies improve. It paints a stark contrast to the dynamic and ever-changing mammalian Y chromosomes that we discuss below.

### Optical mapping assists in chromosomal assembly

The assembly of the chromosome was assisted by the use of optical mapping technology to determine clone order and orientation, and to estimate gap sizes. Optical mapping has been used mainly for small genomes (e.g. Riley et al. 2011), and its use is growing for larger chromosome-level assemblies. Optical mapping assisted with the tomato genome assembly (Shearer et al. 2014), and the recent domestic goat (*Capra hircus*) genome (Dong et al. 2013) was assembled with a combination of short-read sequencing and automated optical mapping that produced super-scaffolds 5 times longer than those generated by conventional fosmid end sequencing. In this case, comparing the previous 10.2 X build with our assembly shows the regions of the chromosome that were particularly helped by the optical mapping approach (see supplementary figure S1). Unsurprisingly, these are the regions enriched for repetitive content; the pericentric heterochromatin especially contained a number of clones in which discrepancies in the sequence could be resolved.

### Improved gene annotation of the porcine X

The revised gene annotation increases the number of protein-coding genes identified on the pig X to 689, bringing the reported gene content closer to that identified in the X chromosomes of well-studied species (i.e. humans and mice, with 813 and 940 protein-coding genes respectively). The majority of X-chromosome genes are shared between species (76% of annotated pig genes shared with human), in accordance with Ohno’s law (Ohno 1967). We have highlighted some specific genes of interest in the Results section with an updated status from build 10.2 X. Beyond this, the improved gene annotation and ordering will be useful not just to the scientific community involved in mammalian sex chromosome evolution, but also to researchers using pigs as a model system for understanding disease. Many loci associated with mental retardation and brain function are found on the human X chromosome (Gécz et al. 2009). We have previously shown an association of X-linked QTLs associated with maternal aggression in sows towards their young with puerperal psychosis in humans (Quilter et al. 2007, 2012); this updated map of the pig X will make detecting significant loci and understanding their contribution to phenotypic traits far more robust in the future. These data also have relevance to the animal breeding industry; many of the QTLs mapped to the X chromosome are associated with production traits (e.g. average daily gain), meat quality traits (e.g. back fat), carcass traits (e.g. percentage lean meat) and reproduction traits (e.g. teat number).

Apart from the olfactory receptor gene clusters, we have not found evidence for widespread ampliconic gene families on the pig X. This contrasts with the X chromosomes of other species; both human and mouse X chromosomes contain independently amplified gene families, with little overlap between the species (Mueller et al. 2013). Human X chromosomes contain multiple inverted repeats with high sequence identity, enriched for testis-expressed genes (Warburton et al. 2004). Mice have a greater number of X-ampliconic genes than humans, apparently driven by a genomic conflict between X and Y sequences; the gene *Sly* on mouse Yq represses gene expression from sex chromosomes in spermatids, and the copy number of X genes has increased in response to maintain expression of key genes (Ellis et al. 2011). These examples led to an expectation that this might be a general feature of mammalian X chromosomes, and that the pig X would also contain unique ampliconic testis-expressed genes, but this appears not to be the case.

### The porcine Y chromosome

One of the striking aspects of the Y chromosome organisation is that the single copy male-specific genes are found in tight clusters of contigs spanning only two or three megabases of sequence. This is a pattern observed in other mammalian Y chromosomes – for example humans (Skaletsky et al. 2003), mice (Soh et al. 2014), cattle (Elsik et al. 2009), cats and dogs (Li et al. 2013a). There is now considerable evidence that each lineage has preserved a small region of key ancestral X genes, and the remainder of the chromosome has evolved in a species-specific manner, often involving expansion of sequences either ancestral or newly acquired into ampliconic tracts.

### Organisation of the pig Y chromsome

#### Palindromic sequences

A recurring feature of the Y sequences we have assembled is the presence of palindromic regions, each on the order of 120kb end to end. These have resulted from duplication and inversion of sequences, and at least three such palindromes are visible. Two have very high levels of sequence identity; the inverted structure may facilitate the process of gene conversion, by allowing the formation of stem-loop structures as seen in the palindromes on the human Y chromosome (e.g. Hallast et al. 2013).

The first of these palindromes is in the proximal gene block, encompassing the two copies of the male-determining gene *SRY*. Multiple *SRY* copies are found in dog (Li et al. 2013a) and some rodent species (e.g. Prokop et al. 2013; Lundrigan and Tucker 1997; Murata et al. 2010), but there has previously been no suggestion of this being the case in the pig. Prior sequencing of the *SRY* gene and comparison across different pig breeds has given no evidence for polymorphisms in the recovered *SRY* sequence from any individual, despite there being breed-specific differences - i.e. there are no heterozygous pigs identified (see >300 sequences deposited in NCBI for *S. scrofa* alone). This suggests that this region undergoes frequent gene conversion that maintains the sequence identity between the copies. While most other species with multiple *SRY* copies have identifiable sequence differences, there is also a known example in rabbits of a palindrome encompassing *SRY*, with gene conversion maintaining sequence identity (Geraldes et al. 2010).

Close to *SRY* lies the second palindromic region, surrounding the *CUL4B* (cullin) fragments. The palindromic region is of similar size to the *SRY* region, and in fact overlaps the *SRY* palindrome. This allowed us to determine that the duplication around *SRY* occurred first, bringing with it the first two cullin exons, after which the cullin fragment was duplicated again (Supplementary Figure S9).

The third example of this phenomenon is found in the distal gene cluster. A palindrome includes the latter 26 exons of the *USP9Y* gene, though without forming an open reading frame. Unlike the previous two palindromes, both breakpoint ends lie within complete transposable elements (TEs). These types of sequence have long been known to facilitate genomic rearrangements via processes including non-allelic recombination and non-homologous end joining (Stankiewicz and Lupski 2002; Hastings et al. 2009). In the case of *USP9Y*, sequence identity between palindrome arms is lower around the breakpoints, perhaps indicating the duplication results from an older event. In all three palindromes, the TEs are annotated as deriving from within the pig lineage – that is, these are not ancient repeat elements. This tells us that these palindromes have arisen relatively recently, and show the ongoing impact of repetitive content in the genome. However, we need data from more species to determine when the duplications occurred. Extant suids diverged since about 40Mya, and the copy number of the genes involved across these lineages remains to be identified.

The palindrome structures are reminiscent of ampliconic structures found on the mouse (Soh et al. 2014) and human Y chromosomes ((Hughes et al. 2012). There are likely other similar palindromic sequences remaining to be discovered within the Y, and more examples of the impact of transposable elements that have promoted accelerated rearrangements will no doubt be found.

#### Ampliconic sequences

Most mapped mammalian Y chromosomes have been found to contain multi-copy gene families (e.g. Li et al. 2013a), and the pig is no exception. Besides the duplications due to palindromes, other sequences have amplified to a much greater extent. There are three gene families of note here.

The *CUL4B* fragments: in addition to the two fragments noted above, further cullin fragments are found proximal to SRY and the active cullin copy, and appear to form part of an ampliconic and fragmented region leading towards the centromere. Cullins are ubiquitin ligase genes, frequently found in multiple copies on mammalian Y chromosomes (Li et al. 2013a; Murphy et al. 2006), and are important for appropriate degradation of substrates during many process including gamete production (Sutovsky 2003).

The *TSPY* fragments: Other amplifications have been defined for *TSPY*, based on FISH mapping of clones. These again appear to be interspersed in the region leading towards the centromere but it not clear how these are arranged. *TSPY* is an ampliconic gene in many mammalian species, from artiodactyls to primates (Xue and Tyler-Smith 2011); the genes are involved spermatogenesis (Xue and Tyler-Smith 2011), and, in cattle, copy number variation of this gene has been linked with fertility in bulls (Hamilton et al. 2012).

The *HSFY* family: These genes are predominantly found in a block between the proximal and distal gene clusters. The organisation and content of this region is complex, and we have investigated it in more detail in a companion paper (Skinner et al, in submission). The *HSFY* genes show evidence for a recent amplification in the *Sus* lineage to ~100 copies, perhaps independent amplification in other suid species, and further independent amplification in cattle (Chang et al. 2013).

All of these pig ampliconic genes are involved in amplifications in other species, and the known functions suggest they are important in spermatogenesis.

#### Other amplified sequences

Yq is dominated by repeat sequences (as demonstrated by the painting pattern of FISH using certain BAC and fosmid clones). These clones are composed almost entirely of sequences related to male-specific repeats described previously for pig (e.g. Mileham et al. 1988), and thus a more detailed study is needed to understand the organisation of this arm of the chromosome. At this time we can say only that patterns of hybridisation from FISH suggest to us that there is single copy sequence on Yq, including near/at the Yq terminus, but we were unable to isolate the sequences involved.

#### Comparative chromosome organisation and gene order between mammals

Previous work from primates, mouse, cat and dog has reconstructed a putative ancestral eutherian Y chromosome (Li et al. 2013a) based on gene order. We have incorporated our pig gene organisation into this, and added information available in the cow Y sequence assembly (Elsik et al. 2009) as shown in supplementary figure S12. One group of genes stands out from the comparison: *USP9Y-DBY-UTY* is the only ancestral cluster of genes that have retained their proximity to each other in all the studied species. There may be a selective disadvantage to disrupting this organisation, as has been proposed for other conserved syntenic blocks in general (Larkin et al. 2009), and for these genes in particular. Both *USP9Y* and *DBY* have been implicated as important in human spermatogenesis, though they may not be essential in all great apes (Tyler-Smith 2008).

*TRAPPC2P-OFD1Y-GPM6B: A potential transposition from the X chromosome* Outside the PAR, there are regions of homologous sequence between the X and Y chromosomes. These can be as small as individual exons of a gene, or multi kilobase regions. We found most of the XY homologies could be attributed to known XY homologous genes, or to otherwise unannotated repetitive sequences, such as ERV families enriched on the sex chromosomes. The large exception was the region *TRAPPC2-OFD1-GPM6B*, which has been subject to recurrent transposition onto the Y from the X chromosome in dog (Li et al. 2013a) and cattle (Chang et al. 2011) lineages. A similar transposition affecting the *RAB9A–SEDL–OFD1Y* genes has occurred in the human and chimp lineage (Hughes et al. 2010). This gene cluster is also at least partially present in the pig Y, with good sequence identity to the X across most of the region. Most discrepancies to the X sequence can be explained by recent transposable element activity (see Supplementary Figure S6 for X-Y alignment and content of the region).

The most likely explanations for the presence of this region in pigs, cattle and dogs are (1) a recurrent transposition of a similar region independently in dogs, pigs and cattle, or (2) in the ancestor of pigs and cattle and independently in dogs. More sequence information in pig is needed to be able to compare the region with cattle to determine if the breakpoints are shared. Our analysis in pig is also complicated by an apparent further rearrangement affecting the region and splitting *GPM6B*, the first exon of *GPM6B* being proximal to *UBE1Y*.

*OFD1* is involved in cilia formation and the ubiquitin-proteosome degradation pathway. Ubiquitin degradation is an important part of gamete development, as seen in the CUL4B genes, and ciliopathies have been previously implicated in fertility issues (Fry et al. 2014). It is plausible that the testis-expressed *OFD1Y* has repeatedly acquired a function in sperm development in different mammalian lineages; certainly the X copies of *OFD1* and also *CUL4B* have been found to be substantially downregulated in teratozoospermic men (Platts et al. 2007).

## Conclusion

This work presents an improvement to the pig X chromosome assembly and gene annotation, plus the first sequence assembly for the pig Y chromosome, produced through a combination of fosmid sequencing, physical mapping by FISH and manual assembly. The assemblies we have generated have allowed new insights into the content and evolution of the pig sex chromosomes, and provide an important resource for the pig genomics community.

## Acknowledgements

We wish to thank Genus for providing the Duroc boar from which a cell line was established and tissue samples obtained. We gratefully acknowledge the Wellcome Trust Sanger Institute core teams for fingerprinting, mapping, archiving, library construction, sequence improvement and sequencing which underpins this work. This work was funded by BBSRC grant BB/F021372/1. The Flow Cytometry and Cytogenetics Core Facilities at the Wellcome Trust Sanger Institute and Sanger investigators are funded by the Wellcome Trust (grant number WT098051). KB, DC-S and JH acknowledge support from the Wellcome Trust (WT095908), the BBSRC (BB/I025506/1) and the European Molecular Biology Laboratory. The research leading to these results has received funding from the European Community’s Seventh Framework Programme (FP7/2007-2013) under grant agreement no 222664 (“Quantomics”). This Publication reflects only the author’s views and the European Community is not liable for any use that may be made of the information contained herein. The authors report no conflict of interests.

## Methods

### Comprehensive methods are given in the supplementary material

#### Library construction and sequencing

Phytohaemagglutinin-stimulated peripheral blood culture from a Duroc boar was used to prepare chromosomes for flow sorting. Flow-sorted Y chromosomes were purified, and 30-50kb sized fragments were cloned into the pCC1Fos vector (library WTSI_1061: http://www.ncbi.nlm.nih.gov/clone/library/genomic/330/). Clones for sequencing were targeted by minimal overlapping clones on a fingerprint contig (FPC) map. The targeted 897 clones for the Y chromosome were sequenced using a combination of 3 different sequencing platforms: Capillary, Illumina and 454. Raw sequence data is submitted to the public data repository, ENA http://www.ebi.ac.uk/ena/, under accession number ERP001277. Clones were assembled using a combination of four assembly scripts to produce *de novo* assemblies. All clone sequences are submitted to ENA. Manual alignment of clone sequences was used to build the clone map, expanding from clones containing known genes. These small contigs were oriented and ordered using fibre-FISH on single DNA-molecule fibres. Final sequence contigs were assembled based on this map using GAP5 (Bonfield and Whitwham 2010).

#### Molecular combing and FISH

Single-molecule DNA fibres were prepared by molecular combing (Michalet et al. 1997). Purified fosmid DNA was amplified using a GenomePlex^^®^^ Whole genome Amplification (WGA) kit, and labelled using a WGA reamplification kit (Sigma-Aldrich) using custom-made dNTP mix as described before (Gribble et al. 2013). Fluorescence *in-situ* hybridisation followed standard protocols. Probes were detected with fluorescently conjugated antibodies. Slides were mounted with SlowFade Gold^^®^^ mounting solution containing 4’,6-diamidino-2-phenylindole (Molecular Probes/Invitrogen) and visualised on a Zeiss AxioImager D1 microscope. Digital image capture and processing were carried out using the SmartCapture software (Digital Scientific UK).

#### X and Y gene annotation, sequence content and chromosomal evolution

Manual annotation on the pig X and Y chromosomes was performed using the Otterlace/Zmap suite of annotation tools (Loveland et al. 2012) following previously established annotation protocols (Harrow et al. 2012; Dawson et al. 2013). The assembled chromosomes were run through an annotation pipeline (Searle et al. 2004), aligning EST, mRNA and protein libraries against the chromosomes with all annotated gene structures (transcripts) supported by at least one form of this transcriptional evidence. The HUGO Gene Nomenclature Committee (HGNC) (Seal et al. 2011) naming convention was used whenever possible for all pig genes, else HAVANA naming conventions (http://www.sanger.ac.uk/research/projects/vertebrategenome/havana) were followed.

RepeatMasker (Smit et al. 1996) was used to identify repetitive elements within Y contigs. Gene content and structure was identified both by gene prediction program GeneMark (http://exon.gatech.edu/GeneMark/) and manual comparison to known Y genes using BLAST. Targeted resequencing was performed across specific genes to confirm their structure (primers given in supplementary table S4). Regions of XY homology were identified by comparing the repeatmasked X assembly to all sequenced repeatmasked Y clones (mapped and unmapped) using LASTZ (Harris 2007). Unannotated repetitive content enriched on the sex chromosomes was minimised by performing an X-X self alignment, and subtracting multiply-hit regions from the X-Y comparison. Candidate regions of 1kb or longer were further analysed using BLAST for similar sequences in the NCBI NT and EST databases, and the VEGA X gene annotations. Evolutionary analyses between X and Y gene pairs were conducted using the Nei-Gojobori model (Nei and Gojobori 1986) in MEGA5 (Tamura et al. 2011). For each pair, positions containing gaps and missing data were eliminated. Reconstruction of ancestral Y chromosome organisations was performed using the Multiple Genomes Rearrangement (MGR) program (Bourque and Pevzner 2002) to calculate optimal rearrangement pathways between each species, as previously described (Skinner and Griffin 2012).

#### Gene expression

RT-PCR was used to confirm expression status of selected genes in five tissues (brain, liver, kidney, side muscle, testis), obtained from the same boar from which blood cultures were derived. Samples were taken from tissues stored in RNAlater (Qiagen) and homogenised in Trizol. Nucleic acids were extracted with phenol-chloroform and DNase treated. RNA was precipitated with isopropanol and stored at 1µg/µl in ddH_2_O at −80°C. RT–PCR was carried out using a OneStep RT–PCR kit (Qiagen) on 25 ng of total RNA. Primer sequences are given in supplementary table S5.

#### Copy number estimation of SRY by qPCR

Primers were designed to amplify a 1447bp region across the majority of the *SRY* ORF and UTRs (F: TAATGGCCGAAAGGAAAGG; R: TGGCTAATCACGGGAACAAC), and products were generated using a MyTaq Red kit (Bioline) using the profile 95°C for 3mins, 35 cycles of 95°C/53°C/72°C for 15s/15s/2min with a final extension of 72°C for 10min. Two female Duroc gDNAs were spiked with dilutions of the purified *SRY* product to give a standard curve of 4 copies *SRY* per genome to 0.25 copies per genome (assuming diploid genome size of 6Gb; Animal Genome Size Database; Gregory 2006). qPCR was performed using a SYPR-FAST qPCR kit (Kapa Biosystems) on the spiked females and on 5 male Duroc gDNAs with primers for SRY and the autosomal (SSC10) gene *NEK7* (supplementary table S6). Annealing temperature was optimised at 57°C. Cycling conditions were 95°C for 3mins, followed by 40 cycles of 95°C/ 57°C/72°C for 10s/20s/30s. The fluorescent signal threshold crossing point (Ct) was normalized to the average signal from Nek7 to produce a normalised ΔC_t_. The data from spiked female gDNA was used to construct a standard curve relating SRY signal to Nek7 signal; from this, an estimate of the absolute *SRY* copy number in the male gDNA samples was produced.

### Data access

All sequence and annotation is available via the Vega genome browser (http://vega.sanger.ac.uk/index.html). The pseudoautosomal region of X/Y homology between the X and Y chromosomes is represented on the X chromosome only in Vega and Ensembl. It is marked as an assembly exception in both chromosomes, but the underlying genomic sequence and annotation is that of the X chromosome. Only the unique regions of the Y chromosome are stored and annotated. The complete Y chromosome is represented by filling the ‘gaps’ with the PAR regions from the X chromosome.

